# A deregulated stress response underlies distinct INF2 associated disease profiles

**DOI:** 10.1101/838086

**Authors:** S. Bayraktar, J. Nehrig, E. Menis, K. Karli, A. Janning, T. Struk, J. Halbritter, U. Michgehl, M.P. Krahn, C.E. Schuberth, H.P. Pavenstädt, R. Wedlich-Söldner

**Author notes:** Equal contributions (S. Bayraktar, J. Nehrig).

## Abstract

Monogenic diseases provide favorable opportunities to elucidate the molecular mechanisms of disease progression and improve medical diagnostics. However, the complex interplay between genetic and environmental factors in disease etiologies makes it difficult to discern the mechanistic links between different alleles of a single locus and their associated pathophysiologies. Here we systematically characterize a large panel (>50) of autosomal dominant mutants of the actin regulator inverted formin 2 (INF2) that have been reported to cause the podocytic kidney disease focal segmental glomerulosclerosis (FSGS) and the neurological disorder Charcot Marie-Tooth disease (CMT). We found that *INF2* mutations lead to deregulated activation of the formin and a constitutive stress response in cultured cells, primary patient cells and *Drosophila* nephrocytes. Using quantitative live-cell imaging we were able to identify distinct subsets of *INF2* variants that exhibit varying degrees of activation. Furthermore, we could clearly distinguish between *INF2* mutations that were linked exclusively to FSGS from those that caused a combination of FSGS and CMT. Our results indicate that cellular profiling of disease-associated mutations can contribute substantially to sequence-based phenotype predictions.

## Introduction

The increasing availability of human genotypes and sequence variation data can yield insights into the genetic basis of many pathological conditions. Monogenic diseases offer a particularly amenable basis for the elucidation of the mechanistic and physiological drivers of disease progression. Focal segmental glomerulosclerosis (FSGS) exhibits several functionally ill-defined pathological phenotypes of glomeruli, the filtration units of the kidney. FSGS is one of the most common types of adult glomerular injury^1^ and has been linked to a combination of genetic and environmental factors^2^. Several autosomal dominant genetic lesions have been linked to FSGS, most of which affect proteins expressed in the podocytes, a specialized renal cell type that forms the epithelial layer of the glomerular filtration barrier. These include genetically determined aberrations in the calcium channel TRPC6, as well as defects in proteins linking the slit diaphragm to the actin cytoskeleton, such as alpha-actinin (ACTN4) and inverted formin 2 (INF2)^3^.

We have previously shown that INF2 globally reorganizes the actin cytoskeleton of mammalian cells in response to calcium influx (Calcium-mediated Actin Reset – CaAR)^4^. CaAR acts as a general stress response that influences diverse physiological processes including transcriptional programs, wound repair and cellular morphogenesis. To date, more than 50 individual mutations in *INF2* have been linked to either FSGS alone or to a combination of FSGS and Charcot-Marie-Tooth disease (FSGS/CMT) (Table 1). Nearly all mutations identified so far map to the N-terminal ‘diaphanous inhibitory domain’ (DID) of INF2, a region that interacts with the ‘diaphanous autoregulatory domain’ (DAD) located in the C-terminal half of the protein, and might thereby contribute to auto-inhibition of the formin^5^. Release of auto-inhibition would also explain the dominant character of *INF2* mutations. Importantly, actin organization in the podocytes of FSGS glomeruli^6, 7^ and in CMT motoneurons^8^ has been shown to be aberrant, and leads to the effacement of foot processes^9^ and disruption of the myelin sheath^10^, respectively.

**Table 1:** Summary of INF2 variants included in this study.

| INF2 variant | Disease | Ref | Polyphen | CADD score | CADD class | cellular phenotype | Actin Reorg / Value |  | Filopodia | CaAR reacting | CaAR max(R) | difference Max(R) to WT |  | difference Max(R) to ΔDID |  |
| --- | --- | --- | --- | --- | --- | --- | --- | --- | --- | --- | --- | --- | --- | --- | --- |
| AA position | no. mutations/ affected |  | score;sensitivity; specificity |  |  |  | +/- | mean ± SD; n | +/- | % cells with Max(R) > 14.9 | Mean ± SD; n | sign. | p-value | sign. | p-value |
| INF2 | control |  |  |  |  | wt | - | 1.03 ± 0.07; 24 | - | 98 | 29 ± 09; 46 |  |  | 1 | 7.44 x 10<sup>-69</sup> |
| ΔDID | control |  |  |  |  | strong | + | 1.21 ± 0.11; 22 | + | 0 | 08 ± 02; 43 | 1 | 7.44 x 10<sup>-69</sup> |  |  |
| A13T | FSGS (1/2) /nat. | 9 | 0.999; 0.14; 0.99 | 28,0 | disease | wt | - | 1.02 ± 0.08; 25 | - | 100 | 28 ± 11; 29 | 0 | 0.419 | 1 | 2.98 x 10<sup>-50</sup> |
| K15R | natural |  | 0.711; 0.86; 0.92 |  |  | wt | - | nd | - | 97 | 29 ± 13; 29 | 0 | 0.006 | 1 | 2.88 x 10<sup>-56</sup> |
| A33T | natural |  | 1.000; 0.00; 1.00 |  |  | impaired | - | 1.03 ± 0.09; 29 | - | 82 | 20 ± 07; 34 | 1 | 6.61 x 10<sup>-13</sup> | 1 | 3.47 x 10<sup>-22</sup> |
| L42P | FSGS (3/3) | 9 | 1.000; 0.00; 1.00 | 24,8 | disease | strong | + | 1.17 ± 0.09; 17 | + | 26 | 12 ± 05; 27 | 1 | 1.36 x 10<sup>-36</sup> | 0 | 0.002 |
| L53P | FSGS/CMT (1/1) |  | 1.000; 0.00; 1.00 |  |  | strong | + | nd | + | 0 | 07 ± 01; 20 | 1 | 1.07 x 10<sup>-46</sup> | 0 | 0.798 |
| L57P | FSGS/CMT (1/1) | 12 | 1.000; 0.00; 1.00 | 26,6 | disease | strong | + | 1.14 ± 0.06; 21 | + | 0 | 08 ± 02; 20 | 1 | 1.81 x 10<sup>-45</sup> | 0 | 0.966 |
| L57R | FSGS/CMT (1/1) | 8 | 1.000; 0.00; 1.00 | 27,5 | disease | strong | + | nd | + | 0 | 08 ± 02; 17 | 1 | 2.08 x 10<sup>-39</sup> | 0 | 0.812 |
| F68S | FSGS/CMT (1/1) | 8 | 1.000; 0.00; 1.00 | 27,1 | disease | strong | + | 1.15 ± 0.08; 24 | + | 4 | 09 ± 06; 26 | 1 | 4.27 x 10<sup>-48</sup> | 0 | 0.413 |
| L69P | FSGS/CMT (2/2) | 39 | 1.000; 0.00; 1.00 | 26,9 | disease | strong | + | nd | + | 0 | 08 ± 02; 19 | 1 | 5.48 x 10<sup>-43</sup> | 0 | 0.890 |
| Δ69-72 | FSGS/CMT (1/1) | 40 |  |  |  | strong | + | nd | + | 0 | 09 ± 02; 24 | 1 | 2.34 x 10<sup>-45</sup> | 0 | 0.373 |
| G73S | FSGS (4/4) | 21 | 1.000; 0.00; 1.00 | 32,0 | disease | strong | + | nd | + | 0 | 08 ± 02; 27 | 1 | 3.57 x 10<sup>-52</sup> | 0 | 0.745 |
| L76P | FSGS (5/5) | 12 | 1.000; 0.00; 1.00 | 26,5 | disease | strong | + | 1.21 ± 0.09; 21 | - | 14 | 10 ± 04; 36 | 1 | 5.97 x 10<sup>-53</sup> | 0 | 0.107 |
| L77P | FSGS/CMT (2/2) | 41 | 1.000; 0.00; 1.00 | 26,2 | disease | strong | + | 1.17 ± 0.08; 22 | + | 0 | 09 ± 01; 20 | 1 | 3.62 x 10<sup>-41</sup> | 0 | 0.469 |
| L77R | FSGS/CMT (2/2) | 40 | 1.000; 0.00; 1.00 | 27,0 | disease | strong | + | nd | + | 0 | 10 ± 02; 19 | 1 | 7.35 x 10<sup>-38</sup> | 0 | 0.279 |
| L81P | FSGS (6/6) | 21 | 1.000; 0.00; 1.00 | 26,0 | disease | strong | + | nd | - | 9 | 09 ± 04; 47 | 1 | 1.21 x 10<sup>-63</sup> | 0 | 0.236 |
| C104F | FSGS/CMT (1/1) | 22 | 1.000; 0.00; 1.00 | 26,4 | disease | strong | + | nd | + | 6 | 09 ± 03; 52 | 1 | 3.23 x 10<sup>-67</sup> | 0 | 0.269 |
| C104R | FSGS/CMT (1/1) | 22 | 1.000; 0.00; 1.00 | 24,6 | disease | strong | + | 1.26 ± 0.12; 22 | + | 3 | 08 ± 04; 34 | 1 | 3.82 x 10<sup>-59</sup> | 0 | 0.728 |
| C104W | FSGS/CMT (3/3) | 22 | 1.000; 0.00; 1.00 | 26,3 | disease | strong | + | nd | + | 0 | 08 ± 02; 26 | 1 | 1.00 x 10<sup>-53</sup> | 0 | 0.907 |
| V105G | FSGS/CMT (1/1) | 42 | 1.000; 0.00; 1.00 | 25,3 | disease | strong | + | nd | + | 0 | 08 ± 03; 30 | 1 | 1.70 x 10<sup>-55</sup> | 0 | 0.756 |
| R106P | FSGS/CMT (1/1) | 22 | 1.000; 0.00; 1.00 | 28,5 | disease | strong | + | nd | + | 0 | 08 ± 02; 26 | 1 | 4.15 x 10<sup>-54</sup> | 0 | 0.858 |
| V108D | FSGS/CMT (1/1) | 39 | 1.000; 0.00; 1.00 | 25,9 | disease | strong | + | nd | + | 5 | 08 ± 03; 22 | 1 | 3.00 x 10<sup>-47</sup> | 0 | 0.899 |
| G114D | FSGS/CMT (2/2) | 40,43 | 1.000; 0.00; 1.00 | 27,3 | disease | strong | + | nd | + | 0 | 08 ± 02; 28 | 1 | 2.93 x 10<sup>-54</sup> | 0 | 0.867 |
| L128P | FSGS/CMT (2/2) | 22 | 1.000; 0.00; 1.00 | 26,7 | disease | strong | + | nd | + | 0 | 08 ± 03; 25 | 1 | 1.23 x 10<sup>-51</sup> | 0 | 0.976 |
| S129P | FSGS (1/1) | 21 | 1.000; 0.00; 1.00 | 23,6 | disease | strong | + | nd | + | 4 | 09 ± 03; 23 | 1 | 6.04 x 10<sup>-44</sup> | 0 | 0.355 |
| L132P | FSGS/CMT (2/2) | 44 | 1.000; 0.00; 1.00 | 26,6 | disease | strong | + | 1.16 ± 0.10; 26 | + | 10 | 10 ± 07; 21 | 1 | 4.57 x 10<sup>-39</sup> | 0 | 0.175 |
| L132R | FSGS/CMT (3/3) | 22 | 1.000; 0.00; 1.00 | 27,5 | disease | strong | + | 1.12 ± 0.06; 20 | + | 7 | 10 ± 04; 27 | 1 | 2.54 x 10<sup>-46</sup> | 0 | 0.190 |
| C151R | FSGS (5/5) | 21,43 | 0.999 ;0.14; 0.99 | 23,8 | disease | strong | + | 1.16 ± 0.08; 19 | + | 4 | 08 ± 03; 26 | 1 | 4.74 x 10<sup>-53</sup> | 0 | 0.995 |
| H158D | FSGS (9/9) | 21 | 0.999 ;0.14; 0.99 | 25,8 | disease | strong | + | 1.16 ± 0.08; 31 | + | 20 | 11 ± 04; 41 | 1 | 7.22 x 10<sup>-49</sup> | 0 | 0.004 |
| H158N | natural |  | 0.997; 0.41; 0.98 |  |  | impaired | - | nd | - | 79 | 21 ± 08; 42 | 1 | 1.67 x 10<sup>-11</sup> | 1 | 1.76 x 10<sup>-28</sup> |

Table1: Summary of INF2 variants included in this study.
| INF2 variant | Disease | Ref | Polyphen | CADD score | CADD class | cellular phenotype | Actin Reorg / Value |  | Filopodia | CaAR reacting<sup>a</sup> | CaAR max(R) | difference Max(R) to WT |  | difference Max(R) to ΔDID |  |
| --- | --- | --- | --- | --- | --- | --- | --- | --- | --- | --- | --- | --- | --- | --- | --- |
| AA position | no. mutations/ affected |  | score;sensitivity; specificity |  |  |  | +/- | mean ± SD; n | +/- | % cells with Max(R) > 14.9 | Mean ± SD; n | sign. | p-value | sign. | p-value |
| L162P | FSGS (6/6) | 19 | 1.000; 0.00; 1.00 | 25,9 | disease | strong | + | 1.16 ± 0.08; 24 | + | 9 | 09 ± 04; 22 | 1 | 8.00 x 10<sup>-42</sup> | 0 | 0.279 |
| L162R | FSGS (3/3) | 42 | 1.000; 0.00; 1.00 | 26,6 | disease | strong | + | nd | ? | 27 | 13 ± 06; 33 | 1 | 1.13 x 10<sup>-38</sup> | 0 | 2.46 x 10<sup>-4</sup> |
| A164P | FSGS (7/7) | 45 | 1.000; 0.00; 1.00 | 28,8 | disease | strong | + | nd | + | 0 | 08 ± 02; 39 | 1 | 1.22 x 10<sup>-63</sup> | 0 | 0.751 |
| L165P | FSGS/CMT (1+1/2) | 22 | 1.000; 0.00; 1.00 | 27,2 | disease | strong | + | nd | + | 4 | 09 ± 02; 25 | 1 | 2.15 x 10<sup>-48</sup> | 0 | 0.567 |
| Δ168 | FSGS (3/3) | 42 |  |  |  | intermediate | + | nd | - | 41 | 15 ± 07; 44 | 1 | 2.19 x 10<sup>-34</sup> | 1 | 5.64 x 10<sup>-9</sup> |
| R177C | FSGS (4/4) | 21,46 | 1.000; 0.00; 1.00 | 34,0 | disease | strong | + | nd | - | 16 | 13 ± 07; 25 | 1 | 2.88 x 10<sup>-31</sup> | 0 | 1.81 x 10<sup>-4</sup> |
| R177H | FSGS (9/11) | 12,43,47, 48 | 1.000; 0.00; 1.00 | 31,0 | disease | strong | + | nd | + | 11 | 11 ± 04; 44 | 1 | 6.93 x 10<sup>-54</sup> | 0 | 0.016 |
| V181G | FSGS/CMT (3+1/4) | 21,49 | 0.998; 0.27; 0.99 | 24,9 | disease | strong | + | 1.19 ± 0.07; 20 | + | 0 | 09 ± 02; 25 | 1 | 1.24 x 10<sup>-48</sup> | 0 | 0.595 |
| E184K | FSGS/CMT (12+3/15) | 8,9,21,42 | 1.000; 0.00; 1.00 | 28,5 | disease | strong | + | 1.18 ± 0.09; 20 | + | 0 | 07 ± 02; 23 | 1 | 7.22 x 10<sup>-51</sup> | 0 | 0.794 |
| E184Q | FSGS (8/8) | 48 | 1.000; 0.00; 1.00 | 25,1 | disease | intermediate | (+) | 1.10 ± 0.09; 20 | - | 87 | 21 ± 07; 30 | 1 | 4.17 x 10<sup>-11</sup> | 1 | 3.89 x 10<sup>-22</sup> |
| S186P | FSGS (18/28) | 9,21 | 0.998; 0.73; 0.96 | 18,5 | polymorphism | intermediate | + | 1.15 ± 0.12; 20 | - | 75 | 17 ± 07; 28 | 1 | 1.17 x 10<sup>-20</sup> | 1 | 2.65 x 10<sup>-11</sup> |
| Y193H | FSGS (1/1) | 12,22 | 1.000; 0.00; 1.00 | 24,1 | disease | intermediate | + | 1.17 ± 0.11; 21 | - | 66 | 18 ± 06; 44 | 1 | 2.58 x 10<sup>-23</sup> | 1 | 3.87 x 10<sup>-16</sup> |
| V194M | natural |  | 0.708; 0.86; 0.92 | 0,9 | polymorphism | wt | - | nd | - | 96 | 26 ± 07; 27 | 0 | 0.026 | 1 | 2.41 x 10<sup>-40</sup> |
| L198R | FSGS (5/5) | 9,12,21 | 1.000; 0.00; 1.00 | 25,3 | disease | strong | + | 1.18 ± 0.11; 31 | + | 4 | 10 ± 02; 26 | 1 | 2.39 x 10<sup>-42</sup> | 0 | 0.068 |
| N202D | FSGS (2/2) | 48 | 1.000; 0.00; 1.00 | 24,9 | disease | strong | + | nd | + | 5 | 09 ± 03; 21 | 1 | 3.19 x 10<sup>-43</sup> | 0 | 0.550 |
| N202S | FSGS (5/5) | 50 | 1.000; 0.00; 1.00 | 23,8 | disease | intermediate | + | 1.19 ± 0.08; 35 | - | 49 | 15 ± 05; 35 | 1 | 5.25 x 10<sup>-31</sup> | 1 | 5.47 x 10<sup>-8</sup> |
| A203D | FSGS (2/2) | 48 | 1.000; 0.00; 1.00 | 27,0 | disease | strong | + | nd | + | 10 | 09 ± 04; 39 | 1 | 6.07 x 10<sup>-61</sup> | 0 | 0.466 |
| R212C | natural |  | 1.000; 0.00; 1.00 | 24,7 | disease | wt | - | nd | - | 88 | 27 ± 11; 26 | 0 | 0.120 | 1 | 5.70 x 10<sup>-43</sup> |
| R212H | natural |  | 0.867; 0.83; 0.93 | 21,6 | polymorphism | wt | - | nd | - | 92 | 24 ± 10; 25 | 0 | 1.14 x 10<sup>-4</sup> | 1 | 2.95 x 10<sup>-30</sup> |
| R214C | FSGS (10/13) | 12,21,42, 48 | 1.000; 0.00; 1.00 | 28,4 | disease | strong | + | 1.13 ± 0.08; 23 | - | 20 | 11 ± 06; 35 | 1 | 8.00 x 10<sup>-45</sup> | 0 | 0.004 |
| R214H | FSGS (20/26) | 9,21,48 | 1.000; 0.00; 1.00 | 27,5 | disease | strong | + | 1.18 ± 0.08; 21 | - | 9 | 10 ± 04; 46 | 1 | 2.42 x 10<sup>-58</sup> | 0 | 0.057 |
| R218Q | FSGS (27/36) | 9,12,21,4 2,48,51 | 1.000; 0.00; 1.00 | 29,5 | disease | strong | + | 1.19 ± 0.08; 27 | + | 0 | 09 ± 02; 27 | 1 | 6.20 x 10<sup>-51</sup> | 0 | 0.596 |
| R218W | FSGS (11/11) | 9,21,42 | 1.000; 0.00; 1.00 | 30,0 | disease | strong | + | 1.24 ± 0.13; 31 | + | 0 | 08 ± 02; 33 | 1 | 1.23 x 10<sup>-58</sup> | 0 | 0.778 |
| E220K | FSGS (14/14) | 9,12,21,4 2,52 | 1.000; 0.00; 1.00 | 27,5 | disease | strong | + | 1.18 ± 0.08; 26 | - | 33 | 13 ± 05; 27 | 1 | 1.43 x 10<sup>-31</sup> | 0 | 4.84 x 10<sup>-5</sup> |
| L245P | FSGS (3/3) | 53 | 0.998; 0.27; 0.99 | 27,9 | disease | strong | + | 1.24 ± 0.13; 24 | + | 0 | 08 ± 02; 24 | 1 | 6.74 x 10<sup>-52</sup> | 0 | 0.827 |
| R261G | natural |  | 0.9998; 0.27; 0.99 | 23,6 | polymorphism | wt | - | nd | - | 89 | 25 ± 09; 27 | 0 | 0.005 | 1 | 2.97 x 10<sup>-37</sup> |
| R261Q | natural |  | 0.929; 0.81; 0.94 | 16,7 | polymorphism | impaired | - | nd | - | 58 | 18 ± 07; 24 | 1 | 4.27 x 10<sup>-16</sup> | 1 | 1.20 x 10<sup>-12</sup> |
| P350L | natural |  | 0.985; 0.74; 0.96 | 23,2 | polymorphism | impaired | - | nd | - | 87 | 22 ± 07; 23 | 1 | 3.43 x 10<sup>-7</sup> | 1 | 6.02 x 10<sup>-23</sup> |
| I685V | FSGS (1/1) / var. | 20 | 0.058; 0.94; 0.84 | nd | nd | wt | - | 1.06 ± 0.06; 23 | - | 91 | 25 ± 08; 22 | 0 | 0.006 | 1 | 2.02 x 10<sup>-32</sup> |
| R877Q | FSGS (1/1) / var. | 20 | 0.113; 0.93; 0.86 | nd | nd | impaired | - | 1.06 ± 0.05; 23 | - | 96 | 23 ± 06; 25 | 1 | 6.44 x 10<sup>-6</sup> | 1 | 2.91 x 10<sup>-27</sup> |

Previous studies have identified a range of interaction partners, posttranslational modifications and potential biological functions of INF2^9, 11–13^. However, while these reports suggest many intriguing links between *INF2* mutations and podocyte pathologies, no systematic analysis of the phenotypic spectrum of known mutations has been performed. Here, we characterize a large panel (>50) of autosomal dominant INF2 mutants that have been reported to cause either FSGS alone or FSGS/CMT (Table 1). We demonstrate that *INF2* mutations lead to deregulated activation of the formin and to constitutive CaAR induction in cell culture, as well as in primary patient cells. Using quantitative live-cell imaging we were able to identify distinct subsets of INF2 variants that mediate different degrees of activation. Furthermore, we could clearly distinguish *INF2* mutations that were associated with isolated FSGS from mutations that were previously reported to cause FSGS/CMT. Finally, we show that expression of INF2 variants in *Drosophila* nephrocytes leads to reorganization of the actin cytoskeleton and altered nephrin distribution *in vivo*, the extent of which is correlated with the severity of the corresponding subset of mutations in humans. Our results indicate that cellular profiling of disease-associated mutations can substantially contribute to sequence-based phenotype predictions.

## Results

### INF2 mediates actin reorganization in primary cells

CaAR is a calcium-dependent cellular actin reorganization process that occurs in cultured mammalian cells, including immortalized human AB8 podocytes. The CaAR stress response is specifically mediated by the formin INF2^4, 14^. Moreover, human osteosarcoma cells (U2OS) expressing disease-linked INF2 variants constitutively exhibit a characteristic reorganization of actin that is reminiscent of that provoked by CaAR^11^. To assess the effects of INF2 mutations on CaAR in physiologically relevant cell types, we first asked whether the stress response can be experimentally induced in primary podocytes. We isolated GFP-labeled podocytes from mouse glomeruli^15^, transiently transfected them with Lifeact-mCherry to monitor actin dynamics, and treated cells with 500 nM ionomycin to induce calcium influx. We observed transient redistribution of actin from the cell cortex to the previously described perinuclear rim^14^ (Figure 1A, Video 1). This phenomenon indeed resembles the CaAR reaction, and occurred with kinetics comparable to those observed in immortalized cells^4^ (Figure 1A). To evaluate the impact of INF2 mutations on cellular actin organization, we then expressed different variants of the ER-localized INF2-CAAX isoform^16^ in isolated primary podocytes. All INF2 variants were N-terminally fused to RFP to enable us to track the subcellular localization of the formin. We initially compared INF2 localization and actin organization for wild-type INF2, a truncated and dominant active form of INF2 (ΔDID) and INF2 point mutants linked to FSGS (L42P) or FSGS/CMT (L57P) (Figure 1B). All INF2 fusions were expressed and localized to reticular tubular networks, as previously described (Figure 1B)^4, 17^. Transient expression of wild-type INF2 did not alter the normal cellular organization of actin, which is dominated by extensive stress-fiber arrays (Figure 1B). Truncation of the autoinhibitory DID region led to a reduction in the incidence of stress fibers and to extensive polymerization of actin in (non-cortical) interior regions of the cells (Figure 1B). This is consistent with the constitutive activity of this mutant reported *in vitro*^18^. Indeed, the disease-linked mutations analyzed here also induced intracellular actin polymerization, most prominently manifested by the formation of a perinuclear actin ring (Figure 1B). Similar actin accumulations have previously been reported for U2OS cells expressing FSGS-linked INF2 variants^11^.

**Figure 1.**
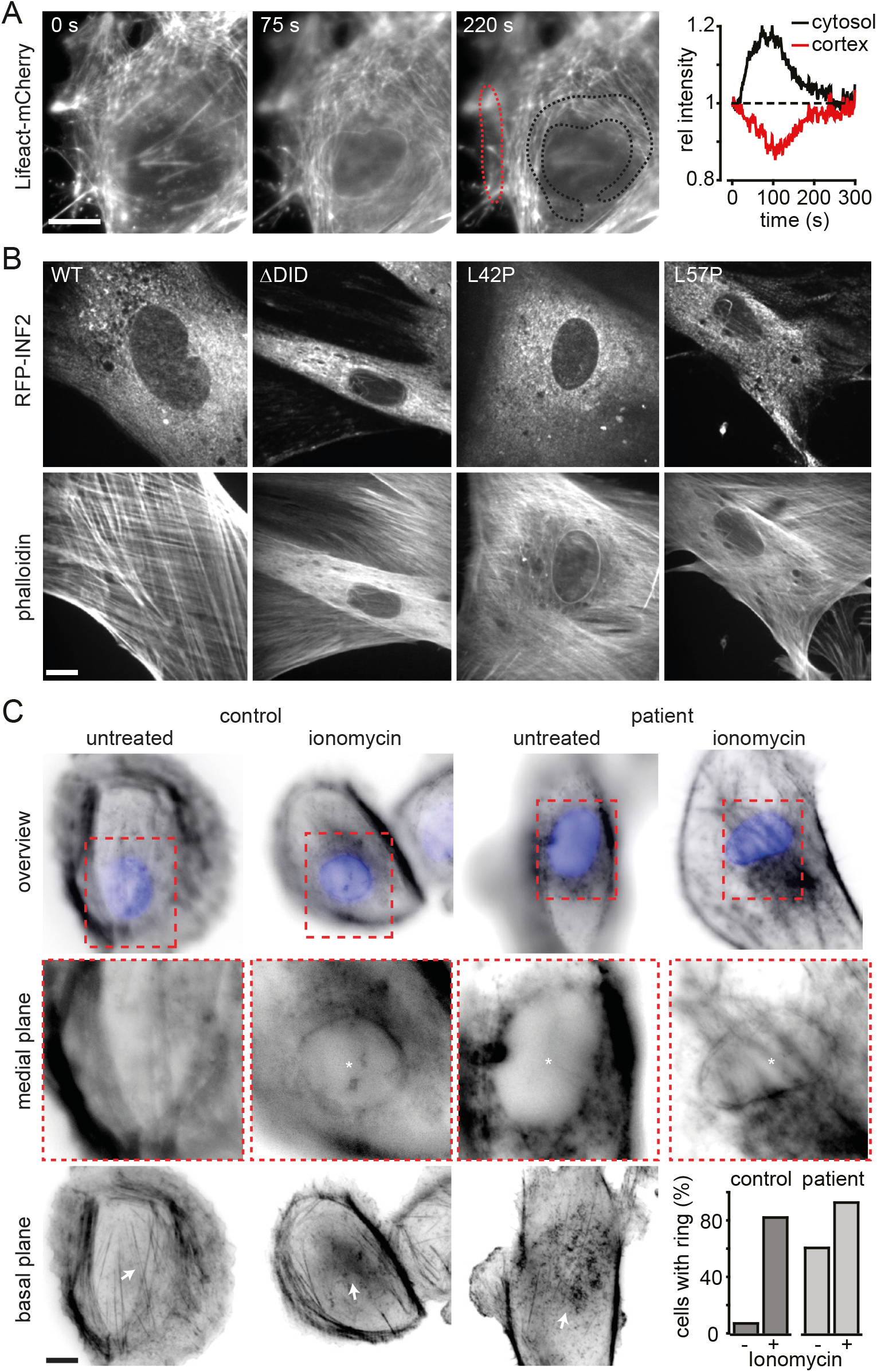
INF2 mediates actin reorganization in primary cells. (**A**) Time-lapse imaging of isolated primary mouse podocytes transfected with Lifeact-mCherry and treated with 500 nM ionomycin at t=0. Dotted lines delineate regions used for intensity plots. (**B**) Images of primary mouse podocytes transfected with indicated RFP-INF2 constructs, PFA-fixed and stained with Alexa350-labelled phalloidin. **(C)** Cells isolated from urine samples obtained from a healthy control and an FSGS patient (INF2 L162P), treated with 500 nM ionomycin for 1 min, PFA-fixed and stained with Alexa594-labelled phalloidin and DAPI. Images were taken at the focal plane of the nucleus. Asterisks indicate cells with intracellular actin polymerization. Arrows indicate changes in actin distribution on the basal surface of the cells (note that fewer stress fibers are present in both ionomycin-treated cells and in patient cells prior to the application of ionomycin). Percentage of cells with intracellular actin is given for n = 15 (control, untreated), n = 28 (control, ionomycin), n = 25 (patient untreated) and n = 15 (patient, ionomycin). Scale bars: 10 µm.

For further validation of these findings, we isolated cells from the urine of a healthy control person and an FSGS patient carrying a dominant *INF2* mutation leading to an L162P substitution^19^. After treatment of control cells with 500 nM ionomycin for 1 min, actin reorganization typical of the CaAR reaction was detected in 82% of the cells examined (Figure 1C). Strikingly, 64% of patient cells exhibited perinuclear actin localization in the absence of ionomycin stimulation (Figure 1C, compared to 6% of control cells). This fraction increased to 91% when patient cells were treated with ionomycin.

Taken together, these results show that INF2-dependent actin reorganization occurs in both primary podocytes and in urine-derived cells from FSGS patients. Disease-linked mutations of INF2 lead to constitutive activation of the formin protein, which results in a stable pattern of intracellular actin accumulation similar to that observed in isolated podocytes after calcium stimulation. Actin reorganization and the CaAR reaction could therefore serve as sensitive and informative readouts for the systematic assessment of disease-linked *INF2* mutations.

### Systematic evaluation of disease-associated INF2 mutations

The majority of FSGS- and FSGS/CMT-related mutations have been mapped to the N-terminal DID of INF2. The only exceptions are two missense variants found in the formin homology 2 (FH2) domain in two sporadic patients, which have not been validated experimentally^20^, and an A13T exchange in the first alpha-helix upstream of the actual DID domain, which was initially linked to the disease but later suggested to be a natural variant^21^. Interestingly, all FSGS/CMT-linked mutations are concentrated in the DID core^22^ (Figure 2A), whereas mutations exclusively linked to FSGS are distributed over the whole domain (Figure 2A). In addition, sequence databases provide an overview of natural variants in INF2 that have not been associated with overt disease. We generated an array of GFP-INF2 fusions, including all known and proposed FSGS/CMT-linked mutations, as well as a representative set of natural variants covering the DID domain (Figure 2A, Table 1). We included wild-type (wt) INF2 and ΔDID as positive and negative controls, respectively.

**Figure 2.**
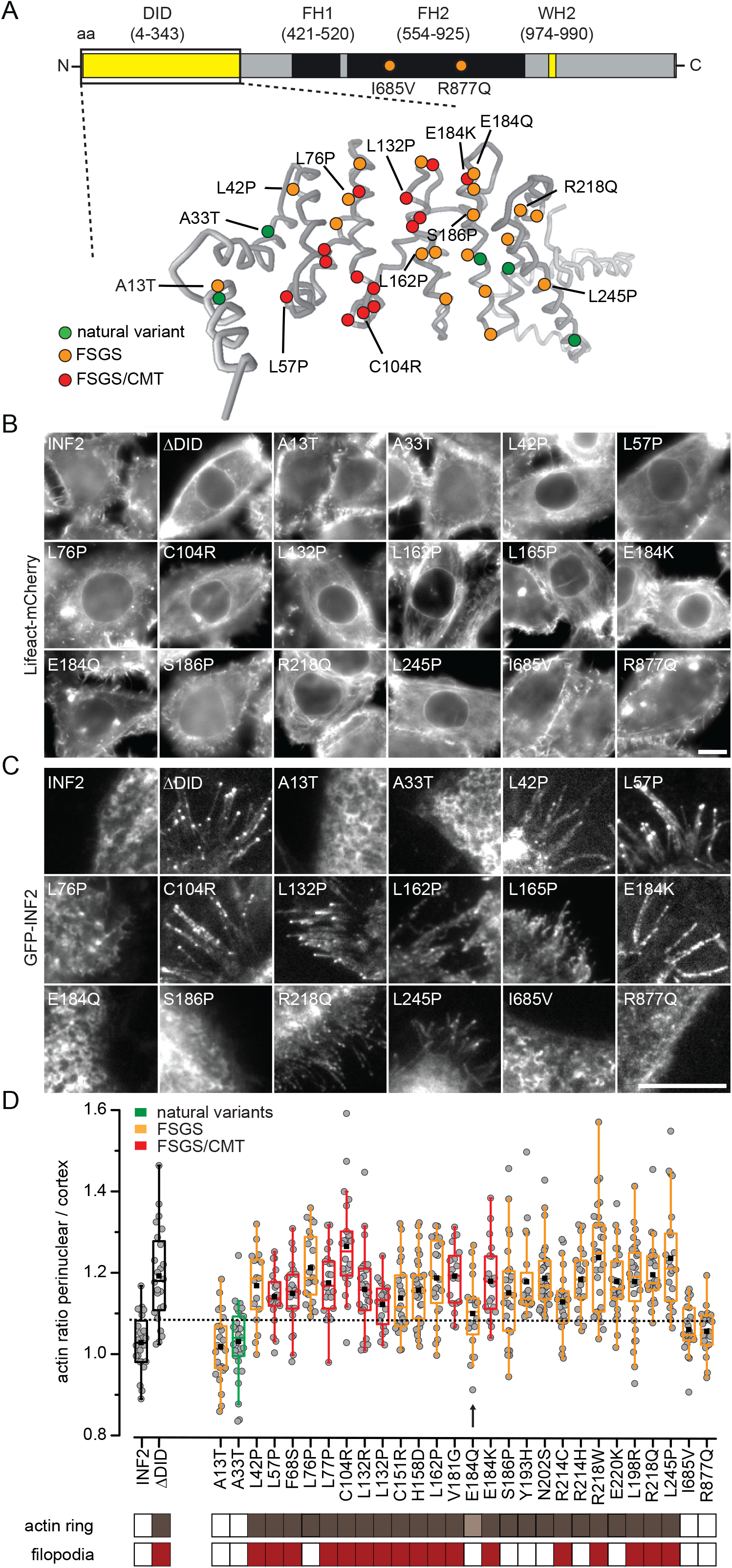
Systematic evaluation of the effects of INF2 mutations on actin organization. **(A)** Overview of INF2 domains and structural model of the INF2 DID domain (amino acids 1 to 343) generated by Phyre^2^. Amino acid substitutions caused by missense mutations are indicated in green, orange and red for natural variants, FSGS-only and FSGS/CMT-linked mutations, respectively. Blue indicates putative FSGS mutations. **(B, C)** Epifluorescence images of HeLA INF2KO cells stably expressing Lifeact-mCherry after transfection with the indicated GFP-tagged INF2 constructs. Images in (C) were acquired by TIRF microscopy to visualize basal GFP-INF2 localization. **(D)** Quantification of perinuclear-to-cortical actin intensity ratios. Cells expressing INF2 wt and INF2 ΔDID were used as positive and negative controls, respectively (black boxes). Natural INF2 variants, FSGS and FSGS/CMT-linked mutations are labelled in green, orange and red, respectively. The dashed line indicates the mean + 2 SD of the wt value. Classification of cells exhibiting intracellular actin rings (brown boxes) and/or filopodia-like structures (red boxes) is shown. Arrow indicates INF2 variant with intermediate actin reorganization values. Scale bars: 10 µm.

We initially tested the effects of INF2 expression on actin organization. We expressed all variants in HeLa INF2KO cells^4^ to rule out spontaneously occurring CaAR reactions mediated by the wt INF2. All of the disease-associated INF2 mutations tested were correctly expressed and localized (Figure 2-figure supplement 1A). Interestingly all variants that lie within the core of the DID domain induced actin polymerization in cells, most prominently displayed as a perinuclear ring (Figure 2B). These effects were observed when variants were expressed in the presence of endogenous INF2 in HeLa wt cells (Figure 2-figure supplement 1B-D). Notably, the naturally occurring INF2 variant A33T, as well as the two I683V and R877Q mutations in the FH2 domain, did not induce prominent intracellular actin accumulation (Figure 2B), while the A13T variant also behaved similarly to wt (Figure 2B).

Upon investigation of the localization of GFP-INF2 in HeLa INF2 KO cells, we observed that, in addition to the ER localization of the INF2-CAAX isoform (Figure 2-figure supplement 1A), active INF2 ΔDID also localized to the tips of cellular protrusions that were filled with actin and resembled filopodia (Figure 2C, figure supplement 2A, B, Video 2). Similar structures were also formed in INF2 ΔDID-expressing primary podocytes (Figure 2-figure supplement 2C) and have been previously reported for HepG2 cells^23^. In agreement with the suggested aberrant activation state of disease-linked INF2 mutants, we found filopodial recruitment of INF2 for many disease variants (Figure 2C, Figure 2-figure supplement 2). For FSGS/CMT mutants, all GFP-INF2 expressing cells scored positive for filopodia, whereas for FSGS-only mutants we found some that did not induce filopodia formation (Figure 2C). Thus the presence of INF2-positive filopodia could provide an initial discrimination between two distinct disease phenotypes.

To evaluate the actin reorganization in more detail, we quantified the actin signal within the transformed cells and compared it to cortical actin levels, as reflected by the actin signal associated with the basal face of the cells. We confirmed that expression of INF2 ΔDID and other disease-associated INF2 variants led to significant increases in intracellular actin filaments (Figure 2D, dashed line corresponds to the 75% quantile). Only the questionable FSGS variant A13T and the two FH2-located mutants resembled the wt distribution. One particular mutation with ambiguous effects on actin reorganization is E184Q (Figure 2D, arrow), which is the most C-terminally located amino acid position to be associated with FSGS/CMT disease. Interestingly, this particular mutation also did not lead to filopodia formation (Figure 2C).

In summary, all *INF2* missense mutations that are clearly associated with disease phenotypes lead to a reorganization of actin within cultured cells, which is consistent with deregulated INF2 activation. This can be monitored both by the CaAR-related accumulation of actin around the nucleus and by the formation of filopodia. The extent of these effects varies, with FSGS/CMT mutations consistently being associated with more pronounced actin reorganization, while mutants exclusively associated with FSGS exhibit a broader range of responses.

### Quantitative kinetic analysis of the functionality of INF2 mutant proteins by CaAR assay

To better discriminate between the effects of different INF2 mutants, we proceeded to establish improved assays for actin reorganization upon INF2 expression. Calcium-mediated actin reorganization during CaAR provides a stereotypical readout for quantitative analysis of INF2 function^3^. To follow actin reorganization live we transfected HeLa INF2 KO cells that stably express Lifeact-mCherry with INF2 mutants and triggered CaAR via laser-induced influx of calcium. We established an automated image analysis routine to quantify the CaAR response in an unbiased manner, using the maximal actin change (max(R)) as the characteristic parameter (Figure 3A, B, Figure 3-figure supplement 1, Video 3). While GFP-INF2-CAAX fully rescued the CaAR reaction in KO cells^3^ (Figure 3A, B), cells expressing the ΔDID mutant showed no reaction to induced calcium influx (Figure 3A, B), which is consistent with constitutive activation of the INF2 mutant, and hence with the stable presence of polymerized actin at the ER and in the perinuclear region. Interestingly, when we expressed FSGS/CMT-associated variants we found two types of responses to CaAR. One subset of mutants, exemplified by Y193H, exhibited actin reorganization upon calcium stimulation, albeit to a lesser extent than wt (Figure 3A, B; Figure 3-figure supplement 1). In others, such as the FSGS/CMT-associated L57P, no further change in actin distribution could be detected, as in the case of INF2 ΔDID expression (Figure 3A, B). To test whether this variable response was a simple consequence of protein levels, we verified comparable levels of expression of INF2 variants by single-cell fluorescence quantification. These results, together with previous reports^13^, indicate that INF2 mutants generally exhibit slightly reduced expression compared to INF2 wt (Figure 3C, Figure 3-figure supplement 2). However, we found no correlation between GFP-INF2 expression and the corresponding max(R) of the CaAR reaction (Figure 3C). This indicates that the magnitude of the CaAR reaction is independent of INF2 levels and can be used for robust quantification of INF2 function (Figure 3D). Furthermore, our results show that the dominant effects of FSGS/CMT-associated INF2 variants can already be discerned at very low expression levels of the mutant protein (Figure 3C).

**Figure 3.**
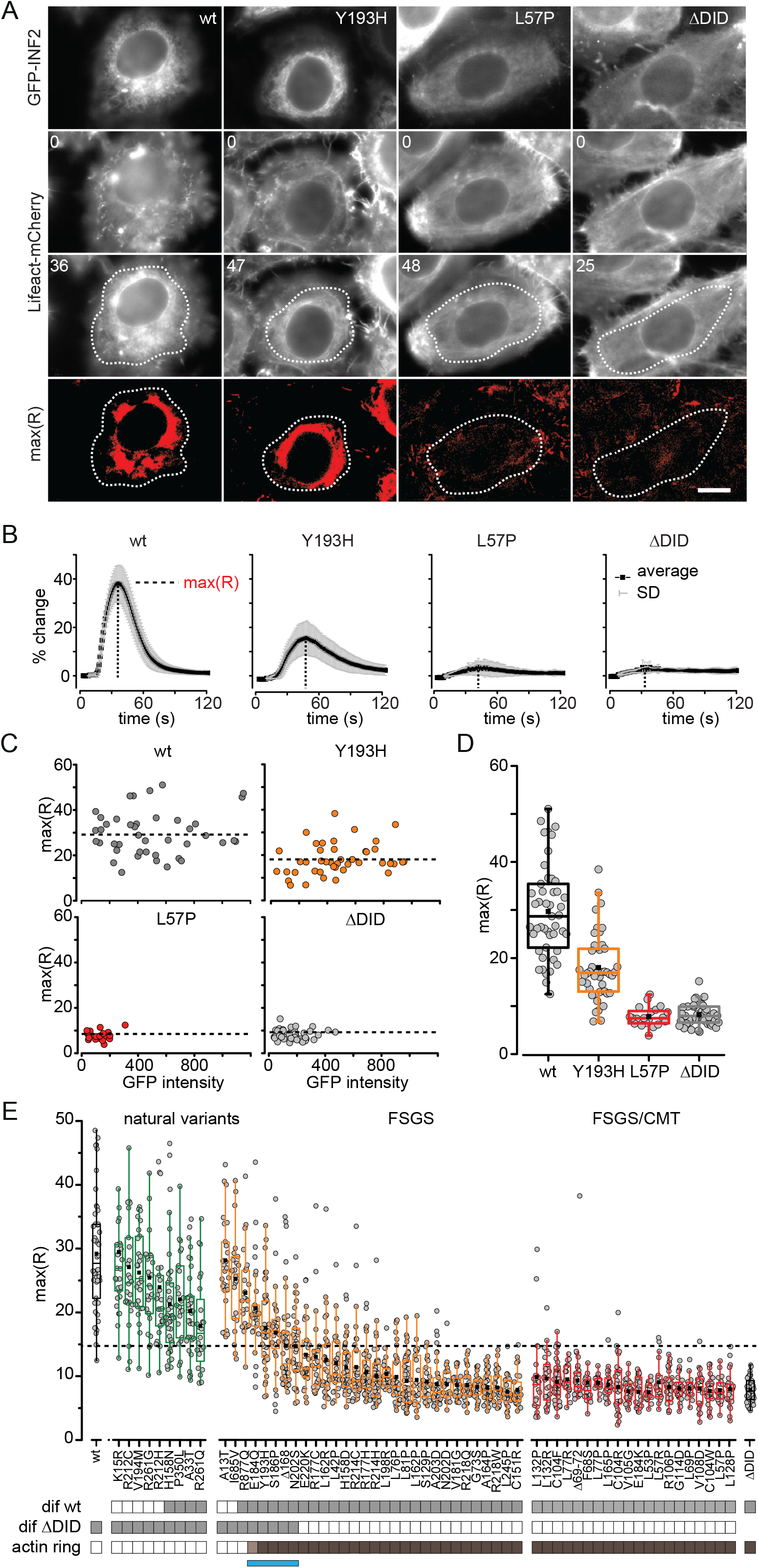
Quantitative analysis of the functionality of INF2 mutant proteins based on the CaAR assay. **(A)** HeLa INF2 KO cells stably expressing Lifeact-mCherry were transfected with the indicated GFP-INF2 constructs. We then monitored actin organization in cells following laser-induced calcium influx at t = 8 s. Images depict actin organization in cells at t = 0 s and at the time point of maximal actin reorganization. Regions used for intensity measurements in (B) are delineated by dashed lines. Scale bar: 10 µm; times are given in seconds. **(B)** Kinetics of actin reorganization observed in (A), given in terms of % change (mean ± SD). **(C)** Graphs of maximal actin reorganization values max(R) for cells expressing the indicated GFP-INF2 constructs, plotted relative to intensity values for GFP-INF2wt. Dashed lines represent means of max(R) values. **(D)** Box plots of max(R) values in (C). **(E)** Box plots of max(R) values for all analyzed INF2 mutations (n ≥ 21). INF2wt and ΔDID were used as positive and negative controls (black boxes), respectively. Natural INF2 variants, FSGS and FSGS&CMT-linked mutations are labelled in green, orange and red, respectively. Values are ordered first by the type of INF2 mutation and then by how much they differ from the ΔDID value (largest to smallest). Grey boxes below the graph indicate that max(R) values were significantly different relative to either wt or ΔDID (one-way ANOVA test with Holm-Bonferroni posthoc correction). Brown boxes indicate GFP-INF2 expression leading to actin reorganization at t=0 s. The blue bar indicates the group of disease-linked INF2 mutations with intermediate actin reorganization phenotypes.

In light of the characteristic max(R) values obtained for different INF2 mutants (Figure 3A-D), we decided to extend our study and comprehensively analyze all reported INF2 mutations. Based on clinical reports and database information ((Figure 2A, Table 1), we separated INF2 variants into three groups: natural variants (green), FSGS-only mutations (orange) and FSGS/CMT-linked mutations (red). All tested natural variants were able to react to calcium stimuli and displayed typical CaAR reactions with averaged max(R) values that were more than 2 standard deviations above the average for ΔDID-expressing cells (dotted line, Figure 3E). In stark contrast, all tested FSGS/CMT variants showed hardly any response to calcium stimulation, with averaged max(R) values similar to ΔDID-expressing cells (Figure 3E). Interestingly, FSGS-only variants displayed a broad distribution of reorganization phenotypes, resulting in max(R) values ranging from wt-like (A13T) to non-responding mutants (C151R). Results for all INF2 mutations examined are summarized in Table 1. To test whether the observed effect of mutant INF2 on actin reorganization also occurs in the presence of wt INF2, we expressed GFP-INF2 variants in HeLa wt cells and measured CaAR kinetics (Figure 3-figure supplement 1). Notably, calcium-induced actin reorganization in the wt background was similar to that observed in KO cells (Figure 3-figure supplement 1).

Statistical analysis (one-way ANOVA, Holm-Bonferroni posthoc test) confirmed that all natural variants differed significantly from INF2 ΔDID. Conversely, all disease-linked INF2 variants, except A13T and I685V, were significantly different from wt INF2 (Figure 3E, grey boxes). We therefore consider the latter two mutations to be natural variants, as suggested in previous publications^6, 7^. Strikingly, we found that a subset of ten intermediate variants resulted in max(R) values that were significantly different from both wt and ΔDID controls (Figure 3E, grey boxes below graphs). Of these, the 4 natural variants and the FH2 variant R877Q did not exhibit any intracellular actin accumulation prior to stimulation. This indicates that these mutations lead to a reduction of INF2 function, rather than deregulated activation. The remaining 5 members of this intermediate group were all FSGS-only mutants and included E184Q, Y193H, S186P, Δ168 and N202S (Figure 3E, blue bar). They exhibited intracellular actin accumulation (Figure 3E, dark brown boxes) independently of the calcium stimulus, with E184Q showing a notably weaker phenotype than the others (Figures 2D, 3E light brown box).

Our results indicate that disease-linked autosomal-dominant INF2 mutations can be divided into distinct subsets using cell-based profiling. To better understand the basis for the observed separation, we set out to further compare molecular interactions and cellular functionalities of the intermediate and fully activated INF2 variants.

### Functional characterization of intermediate mutants

As all disease-relevant mutations are located within the N-terminal portion of INF2, we performed several experiments to study the subcellular localization and molecular interactions of the DID domain (amino acids 1-420). Using yeast two-hybrid (Y2H) assays, we clearly detected interaction between the wt DID domain and the C-terminally located segment of INF2 containing FH1, FH2 and WH2 (Wasp homology 2) domains (FFW, amino acids 421-1008, Figure 4A, B). As predicted from their loss of auto-inhibition, highly active INF2 mutants such as L198R and R218Q showed no interaction with the FFW domain, either as bait or prey (Figure 4A, B). In contrast, intermediate INF2 variants (E184Q, S186P and Y193H) exhibited weak interactions leading to reduced growth on interaction indicator plates (Figure 4A, clearly visible for mutations in the prey) and reduced production of β-Gal (Figure 4B, shown for mutations in the bait).

**Figure 4.**
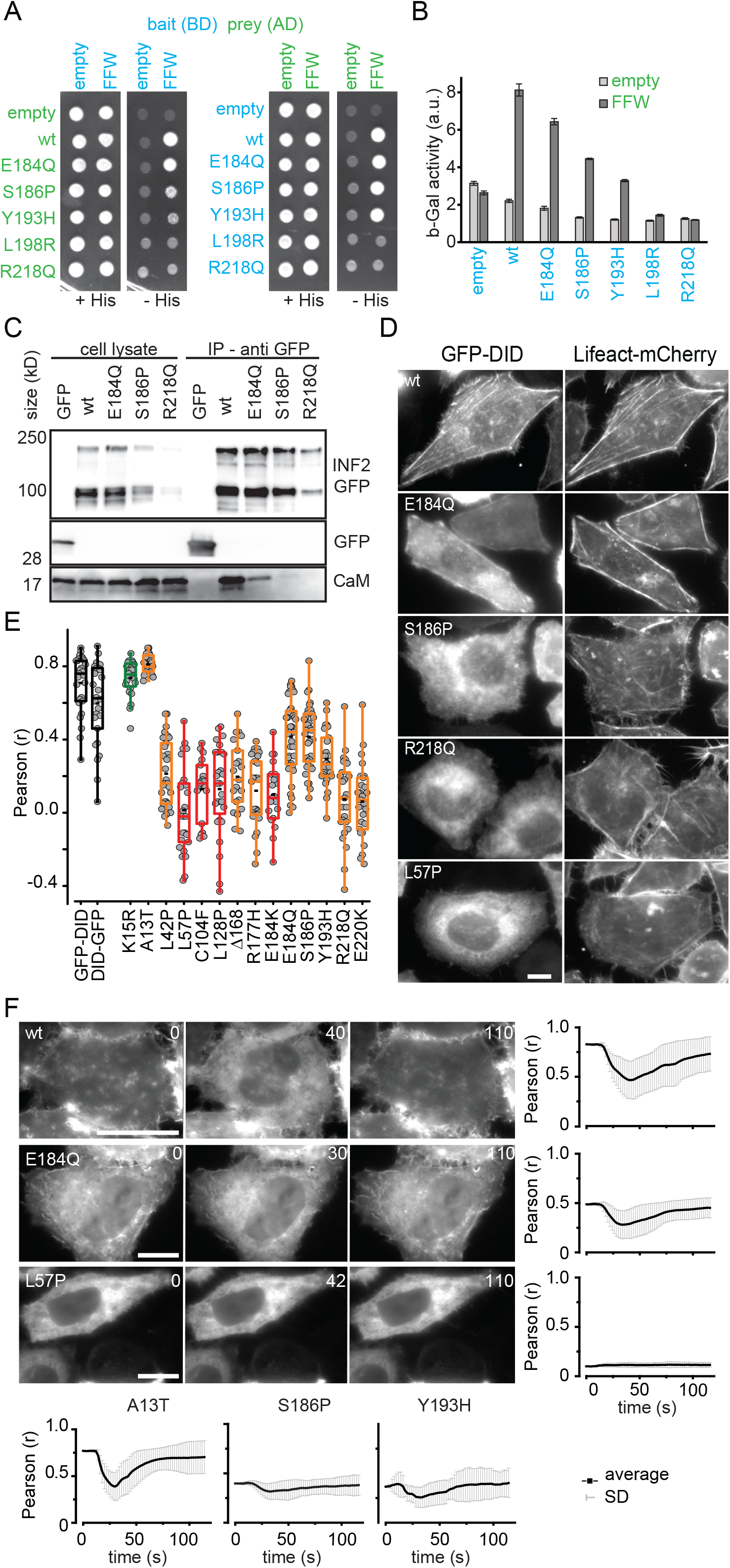
Functional characterization of INF2 mutants of intermediate severity. **(A, B)** Yeast two-hybrid analysis of auto-inhibitory INF2 DID/DAD interaction. Fusions of the INF2 DID (aa 1 to 420) and INF2 FFW (aa 421 to 1008) to the GAL4 activation domain (AD, prey) and the GAL4 binding domain (BD, bait) were expressed in the yeast strain PJ69-4A. The transformed cells were tested for growth on SC-Leu-Trp-His plates (-His) (A) and for beta-galactosidase activity (B), which indicate positive two-hybrid interactions. **(C)** Western blot showing co-IP of calmodulin with GFP-INF2 variants expressed in HEK293 cells. Soluble GFP was used as negative control. Immunoprecipitation was performed using anti-GFP trap. Signal was detected using anti-GFP antibody (INF2-GFP, GFP) and anti-calmodulin antibody (CaM). **(D)** HeLa INF2KO cells stably expressing Lifeact-mCherry were transfected with the indicated GFP-INF2 DID domain constructs (aa 1 to 420) and analyzed for GFP-INF2 DID localization. **(E)** Quantification of colocalization of wildtype (black boxes), natural (green), FSGS-linked only (orange) and FSGS/CMT linked GFP-INF2 DID variants (red) with Lifeact-mCherry was quantified by Pearson correlation coefficient (r) for n ≥ 20 cells. **(F)** Kinetics of GFP-INF2 DID relocalization from actin structures to the cytosol after calcium stimulation, as indicated by the Pearson correlation coefficient (r) of time lapse images with respect to image at t = 0 s. Scale bars: 10 µm.

We have previously shown that INF2 directly binds to recombinant calmodulin^4^. To test whether constitutive activation of INF2 also affected this interaction, we performed co-IP experiments in HEK293 cells. We confirmed that GFP-tagged INF2 wt bound to calmodulin in the presence of calcium (CaM, Figure 4C). This interaction was completely abolished in the case of the strong FSGS mutant R218Q. In contrast, the intermediate mutants S186P and, in particular, E184Q were still able to bind to CaM, but with reduced affinity (Figure 4C).

To test whether the reduced levels of intramolecular interaction measured for mutated DID domains might also affect the subcellular distribution of INF2, we expressed GFP fusions to the DID domain in Hela INF2 KO cells and analyzed their localization. Surprisingly, we found very strong cortical colocalization of GFP-DID with Lifeact-mCherry (Figure 4D, E), which argues for a direct association of the DID domain with actin filaments. This colocalization was completely lost for strong DID mutants (R218Q, L57P, Figure 4D, E). Intermediate DID mutants that retained some ability to interact with the FFW domain in the Y2H assay (Figure 4A, B) also exhibited partial colocalization with actin (S186P, E184Q, Figure 4D, E). The different localization patterns of wt and mutant constructs suggested that the observed actin-binding capacity of the INF2 DID domain might be regulated by calcium in a similar way to the full-length INF2. We therefore stimulated HeLa INF2KO cells by laser ablation to induce calcium signals in the absence of CaAR, and monitored GFP-DID and Lifeact-mCherry signals over time. We observed complete relocalization of the GFP-DID domain from actin structures to the cytosol, which was reversed within 2 min (Figure 4F, Video 4). We used the Pearson correlation coefficient to describe the kinetics of relocalization, which first showed a sharp drop in correlation within 40 s after calcium stimulation, and subsequently returned to the initial level (Figure 4F). When we tested mutant variants of the INF2 DID domain for calcium-induced relocalization, we again found that mutants of the intermediate group were partially able to relocalize, whereas strong DID mutants did not (4F, Video 5). The putative natural variant A13T behaved identically to the wt DID domain (Figure 4F).

In summary, our results suggest that INF2 mutants of the intermediate group retain a reduced capacity for regulation by calcium and mediate stable, albeit weak actin reorganization in cells.

### Physiological consequences of disease-linked INF2 mutations

The findings discussed above show that FSGS- and CMT-linked INF2 variants exhibit disturbed DID domain functionality that leads to partial or complete loss of autoinhibition, altered subcellular targeting of INF2 and constitutive intracellular actin polymerization. We had previously shown that intracellular actin polymerization during CaAR leads to transient inhibition of organelle mobility^4^. To test whether this type of stress response is also induced in cells expressing deregulated disease variants of INF2, we labeled lysosomes in HeLa cells using the live-cell dye LysoTracker Red and then quantified lateral displacements of lysosomes. In HeLa INF2 KO cells expressing wt INF2, lysosomes were highly mobile (Figure 5A). In contrast, in cells expressing the INF2 ΔDID variant, which form stable cytosolic accumulations of actin filaments, lysosomes were found to be concentrated around the perinuclear region (Figure 5A), and exhibited little or no lateral motion (Figure 5A, B). In this respect, all strong INF2 disease variants behaved like the INF2 ΔDID variant, while weaker mutants exhibited intermediate phenotypes with moderately reduced lysosome mobility (Figure 5B).

**Figure 5.**
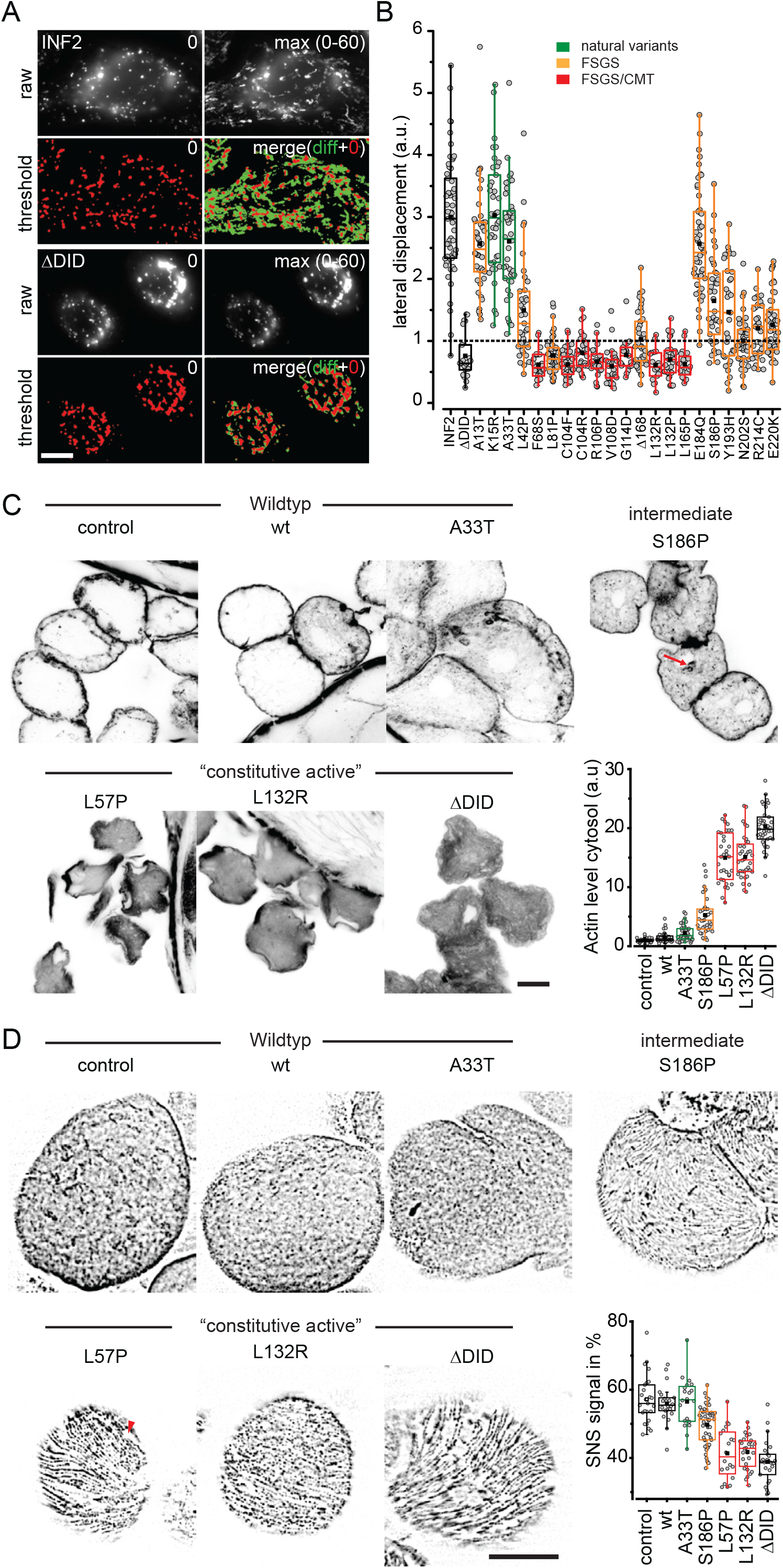
Physiological consequences of disease-linked INf2 mutations. **(A, B)** HeLa INF2 KO cells were transfected with the indicated GFP-INF2 constructs, stained with LysoTracker Red and monitored for the indicated periods. To analyze the distribution of lysosomes in cells, images were thresholded at t = 0 s (red signal) and the area of lysosomal signal at t = 0 s (red in merged image) was compared with the additional area occupied over the period covered by the image series (maximum intensity projection of t = 0 to t = 60 s minus t = 0 s, green in merged image). Box plots in (B) indicate the relative change in the area covered by lysosomes in the course of 60 s, with 1 representing 100% area change (dashed line). Wild-type GFP-INF2 and INF2 ΔDID were used as positive and negative controls (black boxes), respectively. Natural INF2 variants, FSGS and FSGS&CMT-linked mutations are labelled in green, orange and red, respectively. (**C)** Fly nephrocytes stably expressing the indicated Myc-tagged INF2 variants were fixed and stained with phalloidin Alexa647 to analyze actin organization. Graph shows quantification of intracellular versus cortical actin (n≥31 cells). **(D)** Fly nephrocytes stably expressing the indicated Myc-INF2 mutants were fixed and analyzed for nephrin (Sns) localization using anti-Sns antibody. Graph shows quantification of nephrin density (n ≥ 22 cells). Scale bars: 10 µm.

For validation of our results *in vivo*, we expressed different INF2 variants in fly nephrocytes, which are functionally homologous to the podocytes of vertebrates^24^. Using the UAS/Gal4 system, we expressed human INF2 specifically in nephrocytes. Neither fly development nor nephrocyte morphology was visibly affected by INF2 expression (Figure 5C). Alexa647-phalloidin staining of untransfected nephrocytes or nephrocytes expressing wt INF2 showed the expected cortical concentration of actin filaments (Figure 5C). In contrast, expression of the fully activated INF2 mutants ΔDID, L57P and L132R led to strong intracellular accumulation of polymerized actin and reduced cortical staining (Figure 5C). Nephrocytes expressing the intermediate INF2 mutant S186P exhibited punctate intracellular actin staining (Figure 5C, red arrow), but retained continuous cortical actin (Figure 5C). Overall, the ratios of cytosolic to cortical actin correlated well with the suggested levels of INF2 activation in the different mutants (Figure 5C). We next tested whether the altered actin distribution in nephrocytes had any functional consequences. To do so, we monitored the structural organization of slit diaphragms using an antibody specific for the fly nephrin homolog Sns (Sticks and stones). We found the typical dense network organization of Sns in control and INF2 wt-expressing nephrocytes (Figure 5D). Expression of intermediate or strong INF2 variants was associated with reduced Sns staining (Figure 5D) with large areas that were devoid of the protein appearing specifically in strong mutants (Figure 5D, red arrowhead). The discontinuous labeling of membranes with nephrin is comparable to the pattern observed in glomeruli of human FSGS patients^25^.

In summary our results indicate that constitutive redistribution of cytosolic actin driven by disease-linked INF2 variants has wide-ranging consequences for podocyte/nephrocyte structure, including disruption of intracellular trafficking and plasma membrane organization.

## Discussion

We have performed a systematic analysis of actin reorganization in cells expressing INF2 variants. Our study firmly establishes that *INF2* mutations that are linked to familial FSGS and CMT result in aberrant - partial or constitutive - activation of the formin. This activation leads to perinuclear and ER-associated actin accumulation, presumably due to weakening of the autoinhibitory interaction between the mutant DID domain and the DAD domain. This constitutive activity can be observed in primary podocytes and in cells obtained from FSGS patients and is consistent with the observed autosomal dominant nature of these hereditary conditions. In particular, we have shown that several variants linked to FSGS/CMT reduce cortical actin staining (R177H, L165P and R106P^12^, E184K and R218Q^9^) when transiently expressed in cultured cells. These variants were also shown to have increased affinity for constitutively active Cdc42^12^ and decreased affinity for mDia2^26^.

Our evaluation of INF2 mutations using the Calcium-mediated Actin Reset (CaAR) response as a quantitative readout revealed that the degree of INF2 activation varies between different mutant forms of the protein. Mutations associated with FSGS/CMT always resulted in full INF2 activation, as indicated by levels of actin accumulation within cells that were equivalent to those induced by CaAR. In contrast, mutations exclusively associated with FSGS exhibited a broader range of actin responses. Several intermediate INF2 variants, in particular E184Q, Y193H and S186P, retained partial auto-inhibition and were susceptible to further activation by calcium. Interestingly, the S186P variant was reported to retain the ability to bind to mDia2^26^ and not to induce the intracellular actin accumulation associated with other FSGS variants^9^. With our approach we were also able to identify potential false-positive associations of INF2 variants with disease phenotypes. In all our assays, the A13T variant behaved like the wild-type INF2, indicating that the sole reported association of this mutation with FSGS^9^ was erroneous, and confirming the suggestion that it can be classified as a natural polymorphism^21^ that does not compromise protein function. The two FH2 domain variants I685V and R877Q have recently been proposed to be associated with FSGS^20^, but both also behaved like the wild-type control in our cellular assays. Again – each of these variants has only been found in a single patient, and both were considered as potentially false correlations^20^. Our results suggest that CaAR can be used as sensitive cellular assay for INF2 function and for robust evaluation of disease-linked variants of the formin.

During our characterization of molecular interactions of INF2 we surprisingly found that the DID domain on its own closely associated with cortical actin structures. This interaction with actin filaments was sensitive to intracellular calcium levels, and wild-type INF2-DID became completely cytosolic upon calcium influx. Interestingly, a proteolytic N-terminal fragment of INF2 containing the DID domain has recently been shown to associate with the cell cortex^27^, which is consistent with our observation. It is at present unclear how the calcium-regulated actin-binding property of INF2-DID contributes to the function of the full-length protein. However, the DID domain has great potential for use as a rapidly controllable recruitment anchor or marker for actin.

We performed our systematic analyses of INF2 variants in the established HeLa cell line, which is very convenient for genetic manipulations, but it may not reflect the responses of fully differentiated cells types. However, we also validated several key findings in physiologically relevant settings. First, the CaAR reaction, as well as the intracellular actin accumulation upon expression of active INF2 variants, were also seen in cultured primary mouse podocytes and in cells isolated from the urine of human patients. The results obtained with these cells fully confirm the general trend seen in our systematic analysis. Importantly, the degree of intracellular actin accumulation is quantitatively correlated with a perturbed, discontinuous distribution of nephrin in the plasma membrane of fly nephrocytes. This phenotype is reminiscent of other nephropathy models in fly nephrocytes, such as nephrotic syndrome^28^. The defect in nephrin distribution could either be directly linked to a function of cortical actin in the organization of the slit diaphragm^29^ or be due to defects in nephrin trafficking. The latter notion is supported by our findings on the cessation of organelle mobility, which occurs transiently during the CaAR reaction^4^ and is stabilized on expression of constitutively active INF2 variants. A role for INF2 in vesicular trafficking of MAL2 has also been previously proposed for HepG2 cells^23^.

Cellular assays can help to evaluate genetic variants in monogenetic diseases. However, they can only provide a first step towards understanding the mechanistic basis for medical etiologies – in particular for such complex and ill-defined syndromes as FSGS. Many aspects of INF2 regulation have been linked to FSGS, including INF2 expression levels^13^, INF2 interaction with acetylated actin^11^ or MAL^30^, proteolytic processing^27^, or effects on microtubule organization^31, 32^. We favor the idea that INF2-linked FSGS reflects global reorganization of the actin cytoskeleton and thereby a deregulated cellular stress response, rather than specific physiological functions or molecular interactions of INF2 itself. This would be consistent with the aberrant functions of other genetic factors that lead to autosomal dominant FSGS. Thus, FSGS-linked alpha actinin 4 variants have been shown to exhibit calcium-independent bundling of actin, while TRPC6 variants associated with the condition mediate excessive influx of calcium into cells^3^. In all these situations, accumulation of intracellular actin could globally affect intracellular trafficking or the ability of cells to react to acute stress. This interpretation is consistent with the defects seen in transgenic mice that express the strong INF2 FSGS variant R218Q. While kidney development and morphology are not affected in these mice, the ability of podocytes to recover from acute damage induced by protamine sulfate is severely impaired^6^.

## Materials and Methods

### Isolation of primary podocytes

Primary mouse podocytes were isolated from a double-fluorescent-reporter mouse line^15, 33^, in which podocytes are specifically labeled by podocin-Cre-driven expression of GFP, while all other glomerular cells express tdTomato. This mouse line facilitates the separation of podocytes from other glomerular cell types. After cervical dislocation of donor mice, kidneys were removed and washed in Hanks’ Balanced salt solution (HBSS containing calcium chloride and magnesium sulphate, Merck). Kidneys were then cut into small pieces and further dissociated by incubation for 30 min at 37°C in HBSS with 1 mg/ml collagenase (type 2, 255 U/mg, Worthington). To isolate glomeruli, the suspension was then passed consecutively through 100 µm, 70 µm and 40 µm nylon filters (Falcon^™^ Cell Strainer, Corning Glass). The final glomerular fraction was eluted from the filter in 20 ml HBSS, centrifuged at 2000x*g* for 10 min and resuspended in 25 ml RPMI-1640 medium (Merck) supplemented with 10 % FCS (FBS, Gibco), 1% pen/strep (Merck), 1% non-essential amino acids (Gibco) and 1% sodium pyruvate (HyClone). Glomeruli were then cultured for 7 days at 37°C in the presence of 5% CO_2_ in Petri dishes (145/20 mm, Cell Star; Greiner Bio-One) coated with collagen I (Merck). During this time, GFP-positive (podocytes) and Tomato-positive cells grew out of the glomeruli. To sort fluorescent populations, cells were washed with PBS^-2^(Merck), trypsinized and passed through a 40 µm nylon filter (20 ml). The flow-through was centrifuged for 5 min at 1000xg and the pellet was resuspended in 2 ml medium. GFP-expressing cells were sorted by fluorescence-activated cell sorting and subsequently cultivated for 3 days prior to use in experiments.

### Isolation of UREC cells

To isolated urine-derived epithelial cells, total urine was collected, aliquoted into 50-ml tubes and centrifuged at 400xg for 10 min. Pellets were washed once in 20 ml PBS containing 1% pen/strep, and resuspended in DMEM/F12 (Gibco) supplemented with 10% FCS, 1% pen/strep, 1% non-essential amino acids, 1 mM GlutaMax (Gibco), 0.1 mM 2-mercaptoethanol (Gibco) and mixed with REGM^™^ supplements (CC-4127 REGM, Lonza). Cells were seeded into 12-well plates coated with 0.1% gelatin and cultivated for 4 days at 37°C under 5% CO_2_. When confluent (60-80%) colonies became visible, cells were trypsinized for 2 min, centrifuged for 5 min at 400xg, resuspended in fresh medium and transferred to a fresh 6-well plate coated with 0.1% gelatin. Urine-derived cells were cultivated for up to five passages.

### Cell culture and transfection

HeLa cells were grown at 37°C and 5% CO_2_ in Dulbecco’s Modified Eagle medium (DMEM-Glutamax-I; Gibco) supplemented with 10% fetal bovine serum (FBS; Gibco). Routinely, 2×10^4^ cells/ml were seeded into 4-well or 8-well glass-bottomed dishes (Sarstedt) and incubated for 24 h. On the next day, cells were transiently transfected with INF2 constructs using Fugene6 or Lipofectamine 2000 (Invitrogen) according to the manufacturer’s instructions, and incubated for another 24 h. Live-cell imaging was performed with cells in Hanks’ buffered salt solution (HBSS) supplemented with 10 mM HEPES (pH 7.4). The following cell lines were used in this study: HeLa (ECACC 93021013); HeLa cells stably expressing Lifeact-mCherry^34^, and HeLa INF2 KO cells stably expressing Lifeact-mCherry^34^. The identities of all cell lines were checked by visual inspection of morphologies, and all tested negative for mycoplasma.

### DNA constructs and molecular biology

Full length GFP-INF2 constructs were derived from the previously described pGFP-INF2-CAAX vectors that contain siRNA-insensitive coding sequences^4^. The EGFP coding sequence in pGFP-INF2-CAAX was replaced by that of mCherry following digestion with NheI and BglII to obtain pRFP-INF2-CAAX. GFP-DID constructs contain INF2 fragments coding for amino acids 1-420 cloned in the pEGFP-C1 backbone (Invitrogen). All INF2 mutants were generated by site-directed mutagenesis (Stratagene Quickchange, Agilent) using either pGFP-INF2-CAAX or pEGFP-INF2DID as template. For yeast two-hybrid assays, pDEST22 was used as the backbone for prey constructs and pDEST32 for bait. The yeast strain PJ69-4A was co-transformed with both plasmids, selected on SC-Leu-Trp plates and plated onto SC-Leu-Trp-His plates to detect putative interactions. Variants of the INF2 DID domain (aa 1-420, see above) were tested with INF2 FFW (FH1, FH2, WH2; aa 421-1008). Quantitative LacZ analysis was carried out in liquid culture using the ONPG method as described previously^35^.

### Fluorescence microscopy

Epifluorescence imaging was performed on a fully automated iMIC-based microscope (FEI/Till Photonics), equipped with an Olympus 100× 1.4 NA objective and DPSS lasers at 488 nm (Cobolt Calypso, 75 mW) and 561 nm (Cobolt Jive, 150 mW) as light sources. Lasers were selected through an AOTF and directed through a broadband fiber to the microscope. A galvanometer-driven two-axis scan-head was used to adjust laser incidence angles. Images were collected using an Imago-QE Sensicam camera. Acquisition was controlled by LiveAcquisition software (Till Photonics). Ablation experiments were carried out on an iMIC set-up equipped with a pulsed 355 nm picosecond UV laser (Sepia, PicoQuant, Rapp) as previously described^4^. Confocal microscopy was performed on an iMIC42 set-up equipped with a spinning disk unit (Andromeda) using a 60x oil-immersion (NA 1.49) objective. Images were recorded with an EMCCD camera (Andor iXon Ultra 897).

### Immunofluorescence and cell labelling

Primary mouse podocytes and UREC cells were grown on glass coverslips, fixed with 4% paraformaldehyde in PBS for 20 min, washed with PBS and permeabilized with 0.1% Triton X-100 for 10 min prior to incubation with Alexa Fluor 350 phalloidin (Thermo Fisher) or DAPI for 1 h in PBS. The coverslips were washed and subsequently mounted on slides in Mowiol/Dabco (Roth). Lysosomes in HeLa cells were labeled with LysoTracker Red (Thermo Fisher) according to the manufacturer’s directions.

### Image analysis

Images were processed with Fiji prior to data analysis. Linear contrast adjustment and zoom were used for purposes of presentation in Figures only. To measure initial actin reorganization, we acquired z-stacks of cells. For HeLa cell experiments, polygon sections were used as ROIs to define the area around the nucleus and at the basal plane of the cell to measure actin intensities around the nucleus and at the cell cortex, respectively. For nephrocytes, ROIs were selected at the cell outlines for actin intensity measurements at the cell cortex. The initial actin reorganization (a0) was then calculated as: a0 = (a(nucleus) + background)) / ((a(cell cortex)-background).

For analysis of actin reorganization after calcium stimulation we acquired image series at 1 fps. Image series were bleach corrected and the maximum intensity projection of 8 frames prior to stimulation subtracted from all images (reference). We then identified the time point of maximal difference in the series and thresholded this image according to Otsu to generate a binary mask. Max(R) was calculated as relative intensity change (with respect to reference) within the mask at time point of maximal difference. To measure lysosome mobility image series of 60 frames (1 fps) were background corrected (rolling ball, 5 pixel diameter), filtered (median, 2) and thresholded to obtain binary images. Mobility was then calculated as total area covered by lysosomes (diff = max(t_0-60_)-t_0_) relative to the area covered at the start of acquisition: lateral displacement = diff/t0. To quantify colocalization between GFP-DID and Lifeact-RFP images were processed using the Coloc2 plugin of Fiji. Cells of interest were selected by manual ROIs and Pearson correlation coefficients calculated for each time point in a time series. To analyse the density of Sns distribution in fly nephrocytes immunofluorescence images were thresholded and the percentage of total area per cell covered by signal (manually selected ROIs) was determined after background subtraction (rolling ball, 20).

### Statistical analysis

Means and standard deviations are given throughout for quantifications. To test for statistical significance of differences in CaAR max(R) values, one-way ANOVA was performed with the Holm-Bonferroni posthoc correction.

### Characterization of INF2 mutant phenotypes in Drosophila nephrocytes

Fly strains were cultured on standard cornmeal agar food and maintained at 25°C. UASt::Myc-INF2 transgenes were established using the Phi-C31 Integrase system^36^ with *attP40* as landing site. For overexpression of Myc-hINF2, virgin females of the nephrocyte-specific driver line sns::GAL4^37^ were crossed with UASt::Myc-hINF2 males. Garland nephrocytes were isolated from wandering third-instar larvae by dissection, fixed and stained as described previously^38^ using a chicken anti-Sns antibody (1:1000). For phalloidin staining of actin, garland nephrocytes were dissected in PBS pH 7 as described above, and fixed in PBS containing 4% PFA for 10 min. After washing twice with PBS for 3 min each, nephrocytes were permeabilized in PBSTx (0.3% Triton X-100) for 30 min, incubated with phalloidin647 (1:100) in PBS for 2 h and nuclei were then stained with DAPI (diluted 1:1000 in PBS) for 15 min. After two further washes in PBS, nephrocytes were mounted in Mowiol. Images were acquired on a SP8 confocal microscope (Leica).

### Co-immunoprecipitation experiments

Co-immunoprecipitation (CoIP) experiments were performed to detect interaction of GFP-INF2 with calmodulin using the GFP-Trap® affinity resin (Chromotek). All steps were carried out on ice with centrifugations at 4°C. HEK293T cells were seeded on fibronectin-coated (5 μg/ml in PBS) 10-cm^2^ dishes, transfected with INF2 constructs, and harvested by scraping into 1 ml PBS after 24-48 h. After two washing steps with PBS at 500x*g* for 3 min, cells were resuspended in 200 μl lysis buffer (10 mM Tris/Cl pH 7.5; 150 mM NaCl; 500 µM Ca^2+^, protease inhibitors (Roche), 2% NP-40). The suspension was left on ice for 30 min and pipetted up and down several times every 10 min. In the meantime, 25 μl of GFP-Trap® beads were cleared by washing three times in lysis buffer without NP-40. Lysed cells were centrifuged for 10 min at 20,000xg, the supernatant was mixed with 300 μl buffer, and a sample taken for immunoblotting (“total protein”). The rest of the solution was incubated with GFP-Trap® beads by tumbling for 1.5-2 h at 4°C. Afterwards, the suspension was centrifuged at 2500xg for 2 min, washed twice in dilution buffer, and bound protein was eluted from the beads by boiling in 50 μl of two-fold concentrated Laemmli buffer for 5 min. The protein-containing supernatant was analysed by Western blotting.

### Quantification of INF2 expression levels

To analyse expression levels for INF2 variants, HeLa KO cells were seeded in 6-well plates, transfected with GFP-INF2 constructs and grown for 24 h. Cells were then fixed with 4% PFA for 10 min. Nuclei were stained with DAPI, and plates were screened using a ScanR High-Content Screening microscope (Olympus). The object-based autofocus was set on the DAPI channel. For cell selection and automated image analysis, a cut-off was set for nuclear size between 1000 and 5000 pixels. For measurement of the GFP signal, a donut-shaped mask of 20 pixels width was drawn around the nuclear periphery. After subtraction of background signal (14 grey values), mean GFP intensities for all cells in a single experiment (n > 1000) were calculated. The data reported are means of three experiments (N = 3) with the standard error of the mean (SEM).

## Acknowledgements

RWS, HP and CES designed the study and supervised the project. SB, JN, EM, KK, AJ, TS and CES performed experiments and analyzed data. MPK and UM helped with experimental design. JH provided FSGS patient cells. SB, CES and RWS wrote the paper. We thank Hartmut Schmitt for help with isolation of urine cells and Paul Hardy for critical reading of the manuscript. We are indebted to Thomas Zobel and the Münster Imaging Network for help with microscopy. Funding sources: DFG SFB1009-B20 to RWS and HP, IZKF Wed2/022/18 to RWS.

---

**Figure 2-figure supplement 1:**
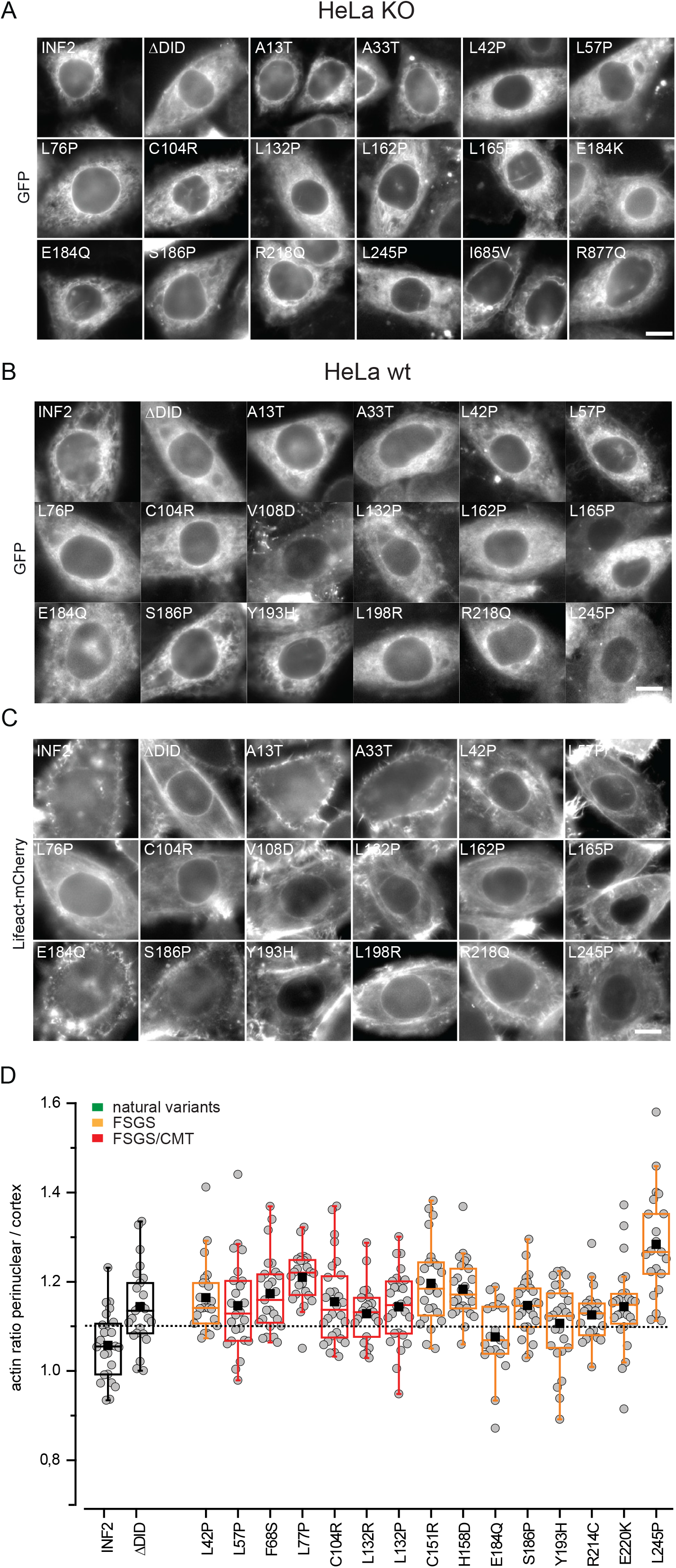
Systematic evaluation of INF2 mutations on actin organization. **(A)** Epifluorescence Images of transiently transfected GFP-INF2 constructs in HeLa INF2KO cells stably expressing Lifeact-mCherry. Corresponds to Figure 2B. **(B, C)** Epifluorescence Images of transiently transfected GFP tagged INF2 constructs (B) in HeLa wt cells stably expressing Lifeact-mCherry (C). **(D)** Quantification of actin intensity ratios of perinuclear over cortical actin in (C). INF2 wt and INF2 ΔDID expressing cells were used as positive and negative controls, respectively (black boxes). FSGS and FSGS/CMT linked mutations are labelled in orange and red, respectively. Dashed line indicates average + 2xSD of wt value. Scale bars: 10 µm.

**Figure 2-figure supplement 2:**
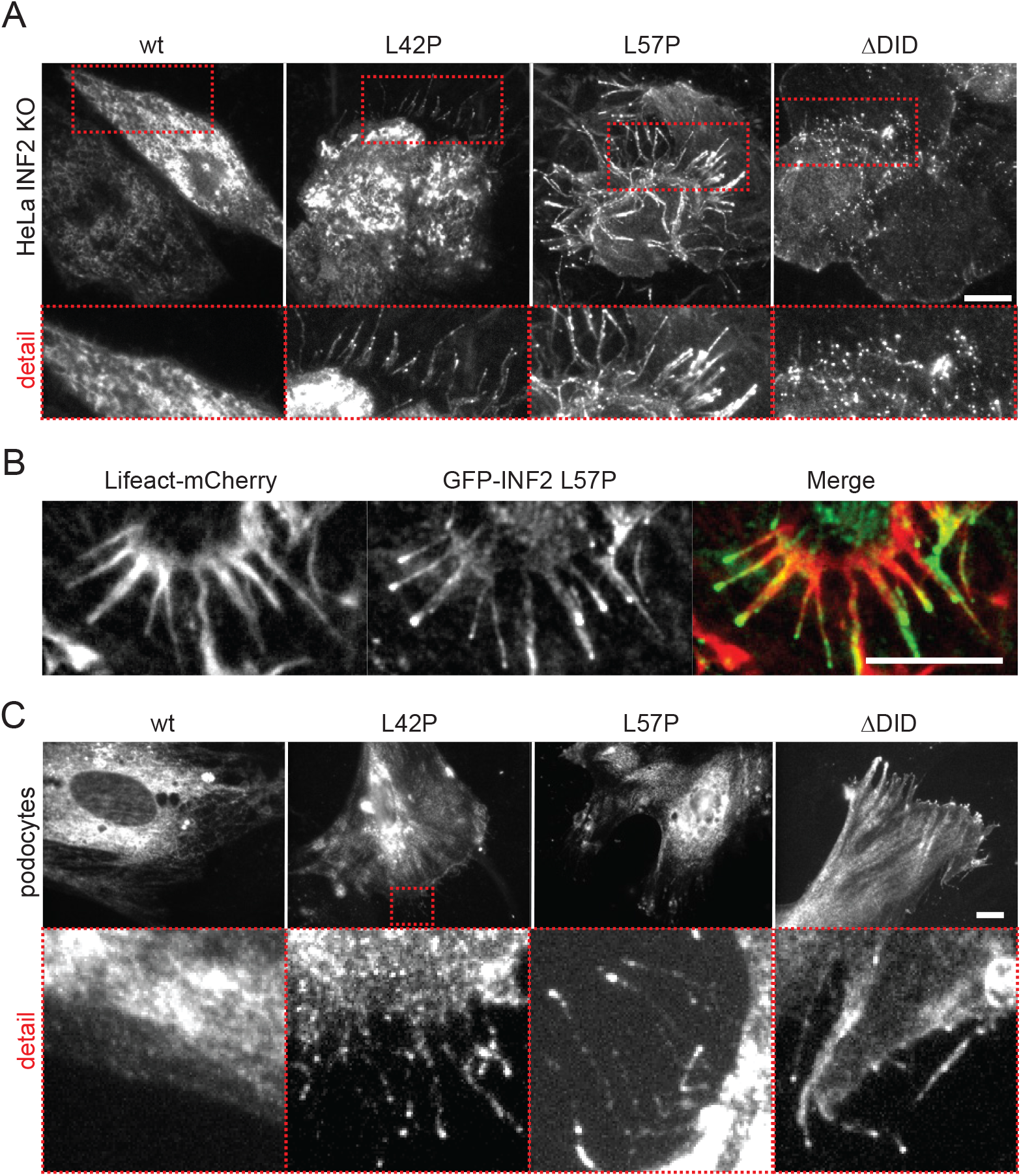
Filopodia-like structures induced by INF2 variants. **(A, B)** TIRFM Images of transiently transfected GFP-INF2 constructs in HeLa INF2 KO cells stably expressing Lifeact-mCherry. Corresponds to Figure 2C. **(C)** TIRFM Images of transiently transfected RFP-INF2 constructs in primary podocytes. Scale bars: 10 µm.

**Figure 3-figure supplement 1:**
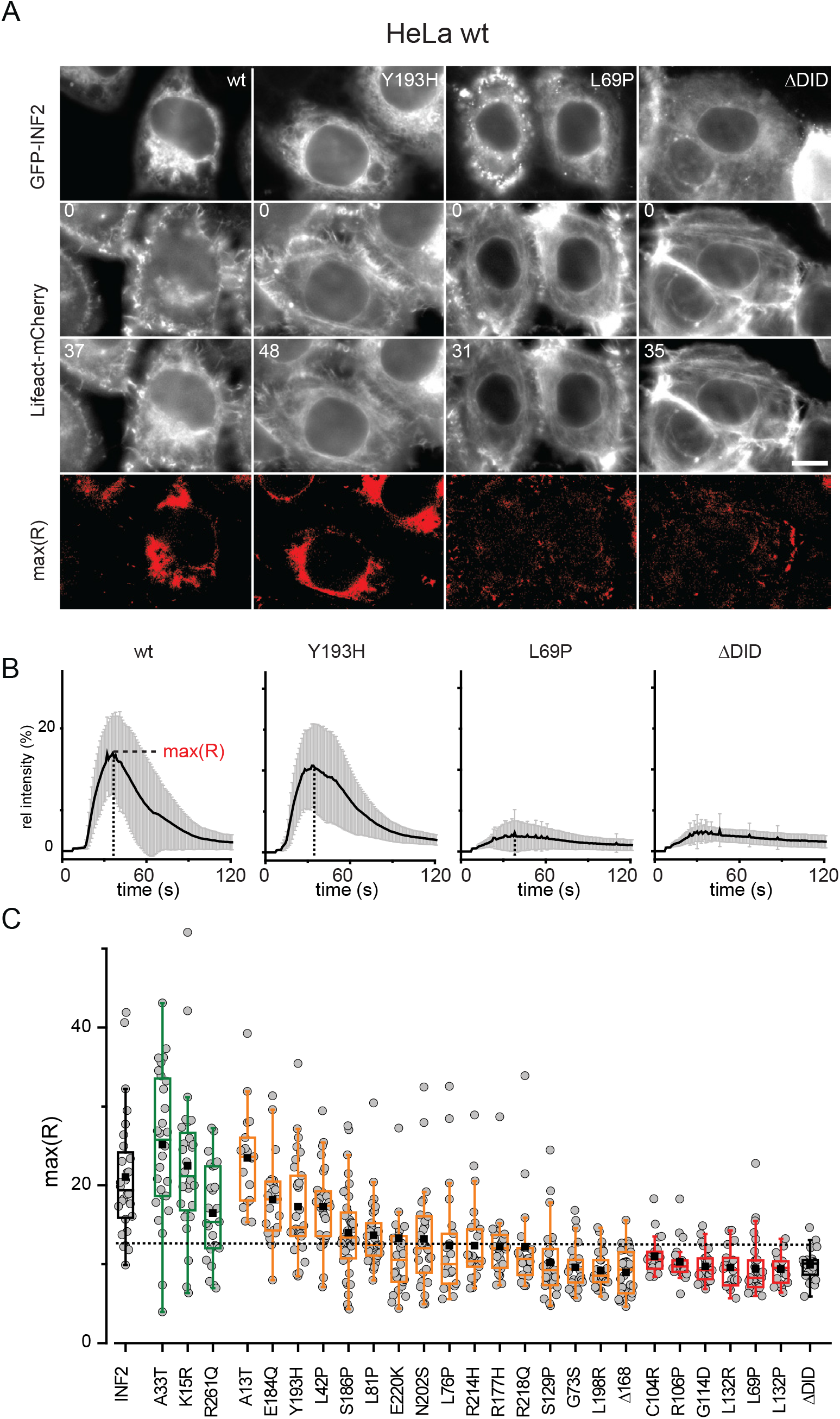
Quantitative analysis of functionality of INF2 mutations by CaAR assay. **(A)** HeLa wt cells stably expressing Lifeact-mCherry were transfected with indicated GFP-INF2 constructs and followed over time with stimulation for calcium influx by laser ablation at t = 8 s. Recordings represented actin organization in cells at the beginning (t = 0 s) and the time point of maximal actin reorganization (indicated time points in seconds and max(R)). **(B)** Kinetics of actin reorganization observed in (A) given as % change (mean ± SD). **(C)** Box plots max(R) values for analyzed INF2 mutations (n ≥ 21). Wildtype and ΔDID were used as positive and negative controls (black boxes), respectively. Natural INF2 variants, FSGS and FSGS/CMT linked mutations are labelled in green, orange and red, respectively. Values are ordered first by type of INF2 mutation and then by difference to ΔDID value (largest to smallest). Scale bar: 10 µm.

**Figure 3-figure supplement 2:**
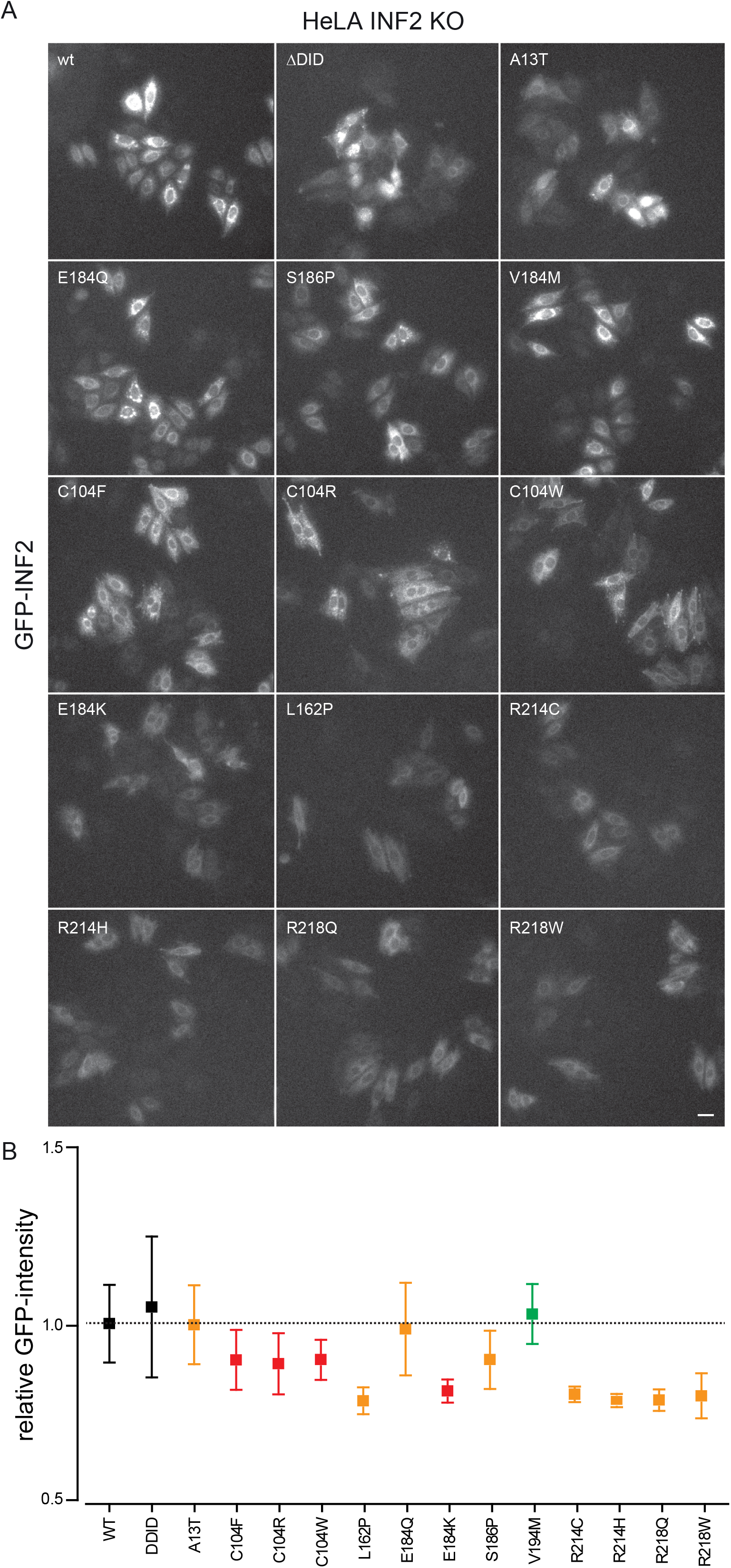
Expression analysis of INF2 variants. **(A)** Epifluorescence images taken on the Olympus ScanR system showing expression of GFP-INF2 variants. **(B)** Quantification of intensities in segmented cells (see methods). Bars indicate Mean ± SEM, n > 500 cells, N = 3 experiments. Scale bars: 10 µm.

**Supplementary file 1. Summary table of INF2 variants included in this study**. Ref: Reference for individual variants; CaAR reacting: % of cells with max(R) value > 14.9, corresponding to mean + 2 SD (ΔDID); sign: Significantly different at significance level 0.01 (1: yes, 0: no); p-values given for ANOVA one-way.

**Video 1.** Primary podocyte transiently expressing Lifeact-mCherry. Cell was stimulated with 500 nM ionomycin at t = 0 s. Corresponds to Figure 1A. Scale bar: 10 µm.

**Video 2.** TIRFM series of HeLa INF2 KO cells transiently expressing indicated GFP-INF2 variants. Corresponds to Figure 2C. Scale bar: 10 µm.

**Video 3.** Epifluorescence series of HeLa INF2 KO cells stably expressing Lifeact-mCherry and transiently expressing indicated GFP-INF2 variants. Cells were stimulated by laser ablation at t = 8 s. Corresponds to Figure 3A. max(R) indicates thresholded image. Time in min:s. Scale bar: 10 µm.

**Video 4.** TIRFM series of HeLa INF2 KO cells transiently expressing GFP-INF2 DID domain. Cells were stimulated by laser ablation at t = 8 s. Associated to Figure 4D. Time in min:s. Scale bar: 10 µm.

**Video 5.** TIRFM series of HeLa INF2 KO cells transiently expressing indicated GFP-INF2 DID domain variants and Lifeact-mCherry. Cells were stimulated by laser ablation at t = 8 s. Associated to Figure 4D. Scale bar: 10 µm.

